# Effects of Age and Cognitive Functions on the Neural Tracking of Speech in Noise

**DOI:** 10.1101/2024.12.26.630452

**Authors:** HyunJung An, JeeWon Lee, Young-jin Park, Myung-Whan Su, Seung-ha Oh, Yoonseob Lim

## Abstract

Older adults often struggle to comprehend speech in noisy environments, a challenge influenced by declines in both auditory processing and cognitive functions. This study examines age-related differences in speech recognition in noise, focusing on the roles of delta (1-4 Hz) and theta (4-8 Hz) neural oscillations and their relationship with cognitive function, particularly working memory. Electroencephalography (EEG) was used to collect data from 23 young adults (20-35 years) and 23 older adults (65-80 years) with normal hearing. Cognitive assessments were administered to older adults, and both groups completed an EEG task involving speech recognition in Speech-Shaped Noise (SSN) at individualized noise levels based on their Sentence Recognition Scores (SRS). Results showed that age significantly impacted hit rates and reaction times in noisy speech recognition tasks. Theta-band neural tracking was notably stronger in older adults, while delta-band tracking showed no age-related difference. Pearson’s correlations indicated significant associations between age-related cognitive decline, reduced hearing sensitivity, and Mini-Mental State Examination (MMSE) scores. Regression analyses showed that theta-band neural tracking at specific SRS levels significantly predicted word list recognition in the higher SRT group, while constructional recall was strongly predicted in the lower SRT group. The findings suggest that older adults may rely on theta-band neural tracking as a compensatory mechanism to support speech perception in noise, with indirect links between working memory and speech perception. Further research is needed to explore the causal relationship between cognitive function and hearing.

## 1. Introduction

Older listeners often experience increased difficulty understanding speech in noisy environments. Advancing age coincides with an observable decline in both peripheral and central auditory processing, which has the potential to affect speech recognition. Previous studies investigating the effect of aging on speech recognition abilities have consistently identified peripheral hearing loss and central auditory processing abilities as major influencing factors, in both quiet and noisy environments (Abel et al., 2000; Souza and Turner, 1994). Age-related damage to the cochlea leads to higher hearing thresholds in the high-frequency range, which is assumed to contribute to difficulties in understanding speech in noise (Gates and Mills, 2005). Decreased speech perception in noise without deficit in peripheral system may be due to a decline in central auditory processing. Central auditory system with reduced grey matter volume in the fronto-temporal cortex (Fjell et al., 2009; Ouda et al., 2015) and diminished white matter microstructure lead to central presbycusis (Wassenaar et al., 2019). Central presbycusis is often indicated by deteriorated auditory object perception and a decline in higher-level auditory processing, including difficulties with speech perception in noisy environments (Alain et al., 2014; Gates and Mills, 2005; Ouda et al., 2015).

Recent studies have attempted to identify the factors contributing to the difficulties of speech perception in noise through neural envelope tracking. Neural envelope tracking was found to be enhanced when focusing on target speech in noise, and this tendency was particularly pronounced in older adults compared to younger adults (Ding et al., 2014; Etard and Reichenbach, 2019; O’Sullivan et al., 2015; Presacco et al., 2019; Vanthornhout et al., 2018). Such enhanced envelope tracking observed in older adults may be due to central compensatory gain mechanisms. These mechanisms increase cortical activity in the brain region to compensate for declined peripheral input. However, it is unclear whether the differences in envelope tracking due to age are a result of basic acoustic processing or attributable to more complex, higher-order factors.

Older adults often face specific perceptual challenges, such as differentiating speech from background noise and recognizing individual words within the stream of acoustic information. They particularly struggle with word recognition when words have many phonetically similar neighbors (Sommers and Danielson, 1999). A possible explanation for these perceptual challenges is that older adults may employ distinct strategies and neural structures to maintain functional speech processing (Tiedt et al., 2020). Studies by Giraud and Poeppel (2012) demonstrated that brain oscillations (specifically delta, theta, and gamma) play a crucial role in speech processing by breaking down continuous speech into discrete units at different timescales that match the natural rhythms of speech (phonemes, syllables, and phrases). Specifically, delta oscillations (1 to 4 Hz) are posited to play an essential role in the segmentation of verbal input into discrete units such as words and phrases, facilitating the integration of lexical units for complex language processing (Ding and Simon, 2012; Schroeder and Lakatos, 2009). Theta oscillations (4 to 8 Hz) facilitated the decoding of acoustic elements (e.g., syllable onset or syllable rate) in speech. Etard and Reichenbach (2019) employed native and non-native speech in various levels of background noise to distinguish between the influence of acoustic attributes and language comprehension. Their findings revealed a primary association of theta-band activity with speech acoustics and delta-band entrainment with comprehension. Moreover, Keshavarzi et al. (2020) revealed that modulating theta-band activity through transcranial alternating current stimulation can influence language comprehension, highlighting the important role of theta frequencies in speech processing. These results align with the notion that the delta band is synchronizing with more complex linguistic elements (words or phrases and other abstract linguistic structures), while theta band oscillations are involved in fundamental acoustic patterns (onset of syllables or syllable rate) (Brodbeck et al., 2018; Ding et al., 2016). Although many recent studies have focused on speech processing mechanisms, only a few studies investigated age-related differences in speech envelope tracking in older adults during speech comprehension (McHaney et al., 2021).

Difficulties in perceiving speech in noise among older adults could be related to lower-level sensory and perceptual changes as well as higher-level cognitive abilities such as working memory span, processing speed, and selective attention (Slade et al., 2020; Fitzhugh et al., 2021). It has been demonstrated that there are connections between cognitive abilities and temporal processing deficits observed during challenging listening tasks (Harris et al., 2010; Schvartz et al., 2008). Moreover, previous studies have shown that the central regions involved in auditory and language processing are critical for speech recognition in noise (Wong et al., 2010; Wong et al., 2008). These findings indicate that central nervous system, which involves cognitive processes, is important for speech recognition in noise.

Among various cognitive processes, working memory is an essential cognitive function for understanding speech in noisy environments (Conway et al., 2001). To perceive speech in noise, a listener rapidly processes incoming auditory information, focuses on the speech, and disregards irrelevant background noise. At the same time, the listener must extract and retain relevant auditory information for integration with subsequent input and later recall. Therefore, it is reasonable to conclude that both working memory and attentional resources are crucial for speech recognition in noise. Specifically, the ability to focus on relevant information while ignoring distractions (selective attention), manage multiple streams of information at once (divided attention), and temporarily store and integrate selected information in working memory are all crucial for perceiving and recognizing speech (Conway et al., 2001; Kane et al., 2001). Several studies have suggested that age-related declines in working memory and speech recognition in noisy environments are significant only for older adults, not younger adults (Fullgrabe and Rosen, 2016; Nagaraj, 2017; Nagaraj and Knapp, 2015). For younger adults with normal hearing, working memory capacity does not significantly predict speech recognition in noise, whereas for older adults, it does. These suggested that working memory and attentional declines in aging may impair speech recognition in noise, underscoring the need to explore these cognitive contributions in depth. Therefore, this study aims to examine how age-related changes in working memory impact speech recognition in noise, addressing a key gap in understanding how these cognitive factors contribute to older adults’ speech processing difficulties in complex auditory environments.

When older adults try to perceive speech in challenging listening environments, they have to rely more on cognitive systems than younger adults. The increased demand on cognitive resources is referred to as listening effort (Classon et al., 2013; McGarrigle et al., 2014). Various subjective and objective methods have been employed to assess listening effort (Peelle, 2018). Subjective measures, including tools such as questionnaires and response times (Gatehouse and Gordon, 1990; Heinrich et al., 2008; Houben et al., 2013), have shown that response times increase under severe noise conditions and that the older group requires more listening effort than the younger group, even in similar listening environments. Conversely, objective assessments of listening effort include techniques like electroencephalography (EEG), neuroimaging, and pupillometry (Kramer et al., 1997; Zekveld and Kramer, 2014; Zekveld et al., 2011). Previous studies showed that while young adults exhibit noticeable increases in pupil dilation during sentence processing correlated with the signal-to-noise ratio (SNR), older adults are less responsive to SNR changes, possibly because they are already utilizing additional cognitive resources for speech understanding in noise (Zekveld et al., 2011). Other studies showed that diminished spectral and temporal resolution in discrimination are known to contribute to the increased demand on cognitive resources during speech understanding in noise (Pichora-Fuller and Souza, 2003; Tun et al., 2009; Ward et al., 2017). Specifically, changes in the amplitude of neural oscillations during speech presentation and memory retention periods measured by EEG have been shown to correlate with variations in listening effort, with an increase in theta band (4–7 Hz) power localized to frontal midline regions reflecting subjective listening effort in speech in noise tasks (Wisniewski et al., 2015). Given these findings, this study aims to investigate how listening effort and working memory demands are reflected in neural tracking during speech processing in noise. By focusing on neural tracking in theta band, this study addresses understanding how listening effort and working memory support speech recognition in challenging listening environments, particularly for older adults. We expect to find that age-related increases in listening effort will correspond with specific patterns in neural oscillations, indicating a greater reliance on cognitive resources for effective speech processing.

This study investigates the neural and cognitive factors influencing speech recognition in noise, with a focus on the effects of age and hearing ability, as measured by Pure Tone Average (PTA) and Speech Reception Threshold (SRT). We first analyzed the role of different cortical oscillation bands, particularly delta and theta, in the neural tracking of speech in noise across various age groups. Additionally, we examined the relationship between cognitive functions—especially working memory—and neural tracking in noise among older adults with different levels of hearing ability. While previous studies have consistently emphasized the impact of peripheral hearing loss and central auditory processing on speech recognition, especially in noisy environments, they have not directly clarified how these age-related auditory declines connect to cognitive function and speech-in-noise perception. To address this, participants were divided into groups based on high and low SRT, we were able to analyze the interplay between cognitive function and speech envelope tracking in noise within each group.

## 2. Material and methods

### 2.1 Participants

Twenty three younger adults (15 males and 8 females; ages 20 to 33 years, mean age = 26 standard deviation = 3.3) who were right-handed with one exception, reported normal hearing and no history of neurological disorders. They enrolled in the study after providing written informed consent.

Twenty three older adults (4 males and 19 females; ages 65 to 80 years, mean age = 72 standard deviation = 3.3) demonstrate that hearing loss was most apparent in the high frequencies and varied from normal to mild hearing loss (HL). To assess the individuals’ hearing ability, we calculated the PTA across the frequency 0.5, 1, 2 and 4 kHz, which ranged from 20 dB hearing level (normal hearing) to 20 - 40 dB hearing level (mild HL). The average hearing threshold was within the normal range even in the older adult group. SRT was determined by the percentage of correctly identified phonetically balanced words from Hahm’s monosyllabic word list. This score represents the dB HL level at which the patient accurately recognizes 50% of the presented words. Older adults were recruited through a screening at Seoul National University Hospital. Adults with no indication of cognitive impairment or learning disability were recruited. The medical history and the presence of learning disabilities were questioned, because serious concussions, medication used to treat, for example, insomnia, and learning disabilities such as dyslexia are known to affect brain responses (Dobreva et al., 2011; Poelmans et al., 2012).

All the procedures in this study followed the ethical standards of the Declaration of Helsinki and were approved by the Institutional Review Board (IRB) of the Seoul National University Hospital (IRB codes:2009-090-1157 and H-2112-075-1282, respectively).

### 2.2 Cognitive testing

The recruited young adults were university students. Given their clear understanding of the oral explanation of the experiment and their ability to follow the procedures without any issues, it was determined that their cognitive levels were within the normal range. Therefore, the cognitive assessment protocols were applied to only older participants to evaluate their cognitive function: Korean Version of Consortium to Establish a Registry for Alzheimer’s Disease Neuropsychological Assessment (CERAD-K) (Lee et al., 2002). The CERAD-K was administered by trained research nurses in conjunction with the psychiatrist’s clinical evaluation. The battery comprises nine tests, listed below in the order they were administered.

1. The Korean version of Mini-Mental Status Examination (MMSE-K): The MMSE (Folstein et al., 1975) is a widely used cognitive screening tool that assesses functions such as orientation, language, attention, visuospatial skills, and memory. In the Korean version, MMSE-KC, the reading and writing tasks were replaced by two judgment-based items to accommodate the significant number of illiterate individuals in Korea. The test has a maximum score of 30.
2. The Korean version of the Short Blessed Test (SBT-K): The SBT-K consists of 6 items that evaluate orientation, delayed memory, and concentration. It has been designed to suit the characteristics of the Korean language. This test includes questions about the current year, month, and time; counting backward from 20 to 1; saying the months of the year in reverse order; and delayed recall of an address and a person’s name. Scores are based on the number of errors in each item, with more errors resulting in a higher score. The maximum score is 28.
3. Word list memory: This free-recall memory test evaluates the ability to learn new verbal information. It involves three trials, each presenting a list of 10 words in a different order. The participant reads each word aloud as it appears. After each trial, the participant has 90 seconds to recall as many words as possible. The Korean version of the Word List Memory task was designed considering Korean language characteristics, including phonemic similarity, semantics, and word length equivalence. The maximum score across the three trials is 30.
4. Word list recall: This test assesses the ability to remember words. Participants have up to 90 seconds to recall the 10 words previously given in the word list memory task. The maximum score for this test is 10.
5. Word list recognition: This test measures the ability to recognize target words from the word list memory task including 10 distractor words. To minimize the chance of guessing correctly, the final score is calculated by adding the total correct answers for both the target and distractor words, then subtracting 10. If the result is less than zero, the score is set to zero. The test has a maximum score of 10.
6. Korean version of the Boston Naming Test (K-BNT): This test assesses visual naming capability by showing 15-line drawings of familiar objects. The drawings are organized into three sets of five, each representing objects with high, medium, or low frequency in the Korean language. The maximum score is 15.
7. Word fluency: This test evaluates verbal production, semantic memory, and language skills. Participants are asked to list as many animals as they can within one minute.
8. Constructional praxis: This task evaluates visuospatial and constructional skills. Participants are asked to copy four-line drawings of increasing complexity: a circle, a diamond, intersecting rectangles, and a cube. Each figure must be copied within 2 minutes. The maximum score for accurately drawing all four figures is 11.
9. Constructional recall: This task evaluates the ability to visuospatial recall, after a brief delay, the four line drawings from the Constructional praxis task. The maximum score for correctly drawing all four figures is 11.

Table 1 presents the demographic characteristics (i.e., age and year of education) and the results of each subtest of the CERAD-K and hearing test.

**Table 1.** Clinical characteristics of old adults.

|  | Older adult |
| --- | --- |
| Age (years) | 72.2 ± 4.6 |
| Educational duration (years) | 11.6 ± 4 |
| CERAD-K |  |
| MMSE-KC | 26.2 ± 1.8 |
| Short Blessed Test | 1.7 ± 1.5 |
| Word list memory | 17.7 ± 3 |
| Word list recall | 71.9 ± 21.1 |
| Word list recognition | 8.5 ± 1.2 |
| Boston naming test | 11.6 ± 1.6 |
| Word fluency | 18.4 ± 3.5 |
| Constructional praxis | 10.6 ± 0.8 |
| Constructional recall | 6.4 ± 2.7 |
| Trail making test A | 54.0 ± 24.7 |
| Trail making test B | 139.5 ± 51.9 |
| Hearing tests |  |
| PTA right ear (dB HL) | 20.4 ± 6 |
| PTA left ear (dB HL) | 19.1 ± 7.5 |
| SRT right ear (dB HL) | 15.8 ± 7.2 |
| SRT left ear (dB HL) | 14.1 ± 6.7 |

\* Pure Tone Average (PTA) ; Speech Recognition Score (SRT) ; Mini-Mental Status Examination (MMSE) ; Korean Version of Consortium to Establish a Registry for Alzheimer's Disease Neuropsychological Assessment (CERAD-K)

### 2.3. EEG experimental design and procedure

#### 2.3.1. Stimuli and experiment procedure

##### Stimuli

In this experiment, the Korean version of the Flemish Matrix Sentence Test was used as the behavioral measure for evaluating the speech recognition of participants, as referenced in a prior study (An et al., 2023). The matrix sentences, consisting of five words representing distinct word categories (name, adjective, object, numeral, and verb), were utilized. To ensure comparable behavioral speech intelligibility scores, 10 alternatives were randomly chosen for each word category, forming a set of sentences with similar characteristics. These matrix sentences were intentionally crafted to natural sound while featuring grammatically simple and semantically unexpected content. This design aimed to minimize the influence of complex language comprehension on the observed results. Additionally, sentences were equated for Root Mean Square (RMS) amplitude at 65 dB Sound Pressure Level (SPL). To explore the speech understanding in noise on neural tracking, matrix sentences were presented under a background noise condition. Speech Shaped Noise (SSN) was employed, representing noise with a spectrum similar to that of the matrix sentences. This allowed for an investigation into how neural tracking is affected by speech understanding in background noise.

##### Experiment Procedure

Before initiating the EEG recording, a speech recognition test was administered to determine the speech understanding level necessary for each individual to accurately recognize speech with the provided speech material (Flemish matrix sentence test). Participants were instructed to repeat each sentence during the test verbally. The adaptive procedure relied on the number of words correctly repeated in each sentence, ranging from zero to five. To determine the noise level, Signal-to-Noise Ratios (SNRs) were established based on participants’ repeated responses. If the participant correctly repeated fewer than 2 words out of the five presented, the noise level was decreased by 3 or 4 dB. However, if they responded correctly with more than 2 words, the noise level was increased by 1 or 2 dB. Through these procedures, we determined the individual SNR required to achieve 25%, 50%, 75%, 95% correct speech recognition scores (SRS). Each participant had an individual SNR level established based on their respective SRS levels.

During the EEG recoding, the participants listened to a 12-minute children’s story titled “Kongjui and Patjui,” narrated by a female Korean speaker. The story was segmented into 1-minute fragments by considering sentence boundaries, thus establishing an optimal condition for constructing a linear decoder, as elaborated in the previous study (An et al., 2023). Following this, the Flemish Matrix sentence test was administered. The participants were instructed to repeat sentences presented under Speech Shaped Noise (SSN) conditions. Across four SRS blocks (SRS 25%, SRS 50%, SRS 75%, and SRS 95%), 10 sentence stimuli were utilized for each SRS block, resulting in a total of 40 sentences. The sentences were presented in random order within each block. To avoid potential order effects, blocks for different sentences were executed in a pseudo-random order. The hit rate was scored as correct if the verbal response was an exact match to the target word. For example, if the target word was “broom”, responses such as “brum” or “broon” were scored as incorrect. The response time was measured from the moment the participants pressed the ENTER key after hearing the sentence, extending to the initiation of their verbal repetition of the sentence heard. The participants were not informed that the response times would be measured to prevent encouraging speed-speech, which could compromise response accuracy and induce speed-accuracy trade-offs (Heitz, 2014).

**Figure 1.**
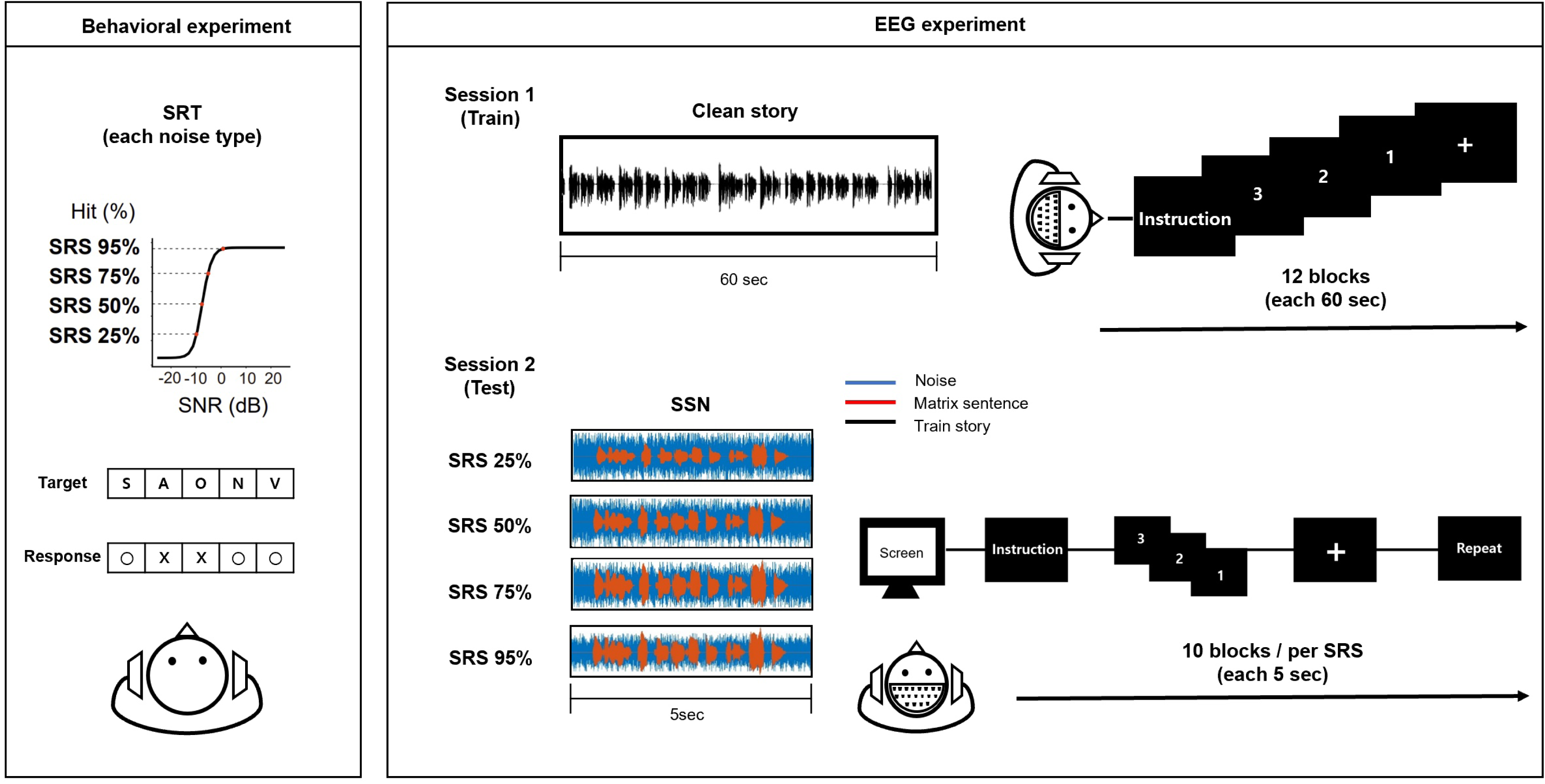
Overview of the main procedures conducted during the behavioral and EEG experiment. In the behavioral experiment, the participants were directed to repeat each sentence heard verbally. The noise level was adjusted based on the number of words correctly repeated in each sentence, which varied from zero to five. For the EEG study, noise levels were selected to match specific target speech recognition levels (SRS: 25%, 50%, 75%, and 95%). The EEG experiment procedure was structured into two sessions. In the first session, a decoder model for speech recognition was developed using the narrative “Kongjui and Patjui” without noise. In the second session, SSN was presented at varying levels of SRS (25%, 50%, 75%, and 95%), as determined by the behavioral test. For each level of noise, the participants were presented with 10 sentence stimuli within four blocks of noise conditions. They were required to attend to each sentence under noise conditions and repeat what they heard within a limited time (8 sec).

#### 2.3.2. Acquisition and pre-processing of EEG data

The EEG experiment was conducted in an electromagnetic-shielded, double-wall, soundproof chamber that meets the ambient noise level requirements of the American National Standards Institute (ANSI) S3.1-1999. EEG data were collected using a 64-electrode Neuroscan SynAmps RT system (64-channel Quik-Cap from Compumedics, Victoria, Australia) at a sampling rate of 1000 Hz and raw signals were re-referenced through a common average reference method, excluding vertical and horizontal electrooculograms.

The preprocessing of all data analysis was conducted using EEGLAB and the mTRF toolbox (Crosse et al., 2016) in MATLAB (v9.5.0 R2018b, The MathWorks, Inc., Natick, MA, USA). The preprocessing procedure involved applying bandpass filtering to the original recordings using a zero-phase Hamming windowed sinc FIR filter in the delta range (1–4 Hz) and theta range (4–8 Hz). After filtering, both the EEG and speech envelope data were downsampled to a sampling rate of 64 Hz to maintain uniform sample lengths across datasets. Additionally, z-scoring was employed to normalize the data.

We analyzed the decoding by distinguishing the neural activity in the cerebral cortex to understand the mechanism of speech in noise conditions.

#### 2.3.3. Estimation of neural tracking of speech

To examine how speech perception processes and neural representations of speech are affected under noisy conditions, we employed a stimulus reconstruction method as described by Vanthornhout et al. (2018). In this method, a speech envelope is reconstructed from EEG signals recorded while participants listen to speech stimuli. The degree of similarity between the original and reconstructed speech envelopes indicates the accuracy of the cortical representation of actual speech and can vary depending on the listener’s level of attention(Ding and Simon, 2014; Vanthornhout et al., 2018; Calderone et al., 2014). Based on this method, we calculated the linear decoder for the EEG signals to generate a reconstructed envelope of the stimulus, denoted as *Ŝ*(*t*). The mathematical representation of this reconstruction is as follows:

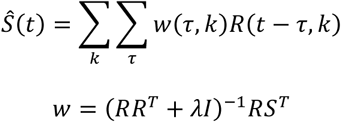

*Ŝ*(.) denotes the reconstructed envelope, where *t* corresponds to the time range, *k* refers to the index of the recording electrodes, τ represents the time lag between the stimulus and the neural response, and R(.) indicates the neural response. To determine if a linear relationship between the EEG data and the envelope of speech stimuli, ridge regression was applied using *w*(.) as weights of decoder.

A subject-specific decoder was first calculated for the EEG recordings of each participant while listening to the “Kongjui and Patjui” story. Then, the subject-specific decoder was applied to the EEG recordings obtained during the Flemish Matrix sentence test under different SNR levels, which estimates the envelope of the clean matrix sentences. The correlation between the estimated envelope and the envelope of the clean matrix sentence was assessed using Pearson’s correlation coefficient. The time-lag parameter τ was selected within a range of 0-500 ms, following guidelines from previous studies (O’Sullivan et al., 2015). The L2 regularization parameter λ was set at 10^^3^ determined through an iterative search process aimed at optimizing detection accuracy (Wong et al., 2018).

### 2.4 Statistical analysis

Statistical analysis was performed using the R (version 3.3.2) software. A p-value of 0.05 was considered to indicate statistical significance.

Initially, we employed a two-way mixed analysis of variance (ANOVA) to assess the effect of varying noise levels on behavioral performance, including the hit rate and reaction time. The hit rate and reaction time data underwent independent analyses using a 2 (age group: Younger subjects, Older subjects) × 4 (SRS condition: 25%, 50%, 75%, and 95%) mixed ANOVA. Additionally, ANOVA analyses were conducted on the neural tracking to explore how different frequency bands (delta and theta) varied with noise levels based on age group. This methodological approach enhanced our understanding of the interaction between age and noise levels in both the behavioral and neural measurements. The F scores and p-values derived from the ANOVA analysis were utilized to ascertain the statistical significance of the observed difference. A further exploration of specific group comparisons and identification of significant pairwise differences were conducted through post hoc model comparisons using Bonferroni-corrected paired t-tests.

Secondly, to explore the relationship between cognitive abilities and speech understanding or age, we conducted a descriptive statistical analysis of the demographic data, hearing tests, cognition tests, and language assessments. Pearson correlation analysis was conducted to examine the relationships between the participants’ age, SRT, PTA, neural tracking of SRS in noise, MMSE, and CERAD-K test. The CERAD-K consists of two parts: 1) verbal working memory (word list memory, word list recall, word list recognition), and 2) visual working memory (constructional recall). In this analysis, we focused on visual and verbal working memory, as these areas are closely related to the cognitive domains involved in speech recognition in noise.

Lastly, the multiple regression analysis was performed with the working memory task results as the dependent variable and the neural tracking of SRS at different noise levels (25%, 50%, 75%, and 95%) as the independent variables (working memory part of CERAD-K). Finally, to explore the relationship between working memory and neural tracking of SRS at noise levels according to SRT, a multiple regression analysis was conducted, dividing the participants into higher and lower groups based on the median value of their SRT. Statistical significance was set at p < 0.05.

Throughout all of our analyses, we calculated effect sizes to evaluate the strength of our results. Specifically, an eta square effect size (η^2^) was categorized as follows: 0.01 indicates a small effect, 0.06 a medium effect, and 0.14 a large effect. Similarly, for Cohen’s d, effect sizes will be interpreted with 0.2 representing a small effect, 0.5 a medium effect, and 0.8 a large effect (Lakens, 2013).

## 3. Results

### 3.1 Behavioral evaluation of SRS

To compare listening effort in terms of accuracy and reaction time, we conducted a two-way mixed ANOVA involving age groups (2) and SRS conditions (4). Figure 2A revealed a significant main effect of the SRS conditions on hit rate (F_(3, 132)_ = 107.432, *p* = 0.0002, η^2^ = 0.39), indicating a notable influence of varied SRS noise conditions on the participants’ hit rate. However, there was no significant main effect of age group (F_(1, 44)_ = 3.343, *p* = 0.0743), suggesting that age did not significantly affect the hit rate. The interaction between age group and the SRS noise conditions was not significant (F_(3, 132)_ = 1.795, *p* = 0.151), implying that, as noise levels decreased, the hit rate increased for both age groups. Post-hoc comparisons revealed significant differences in hit rates, with all SRS levels showing significant differences (p < 0.05).

**Figure 2.**
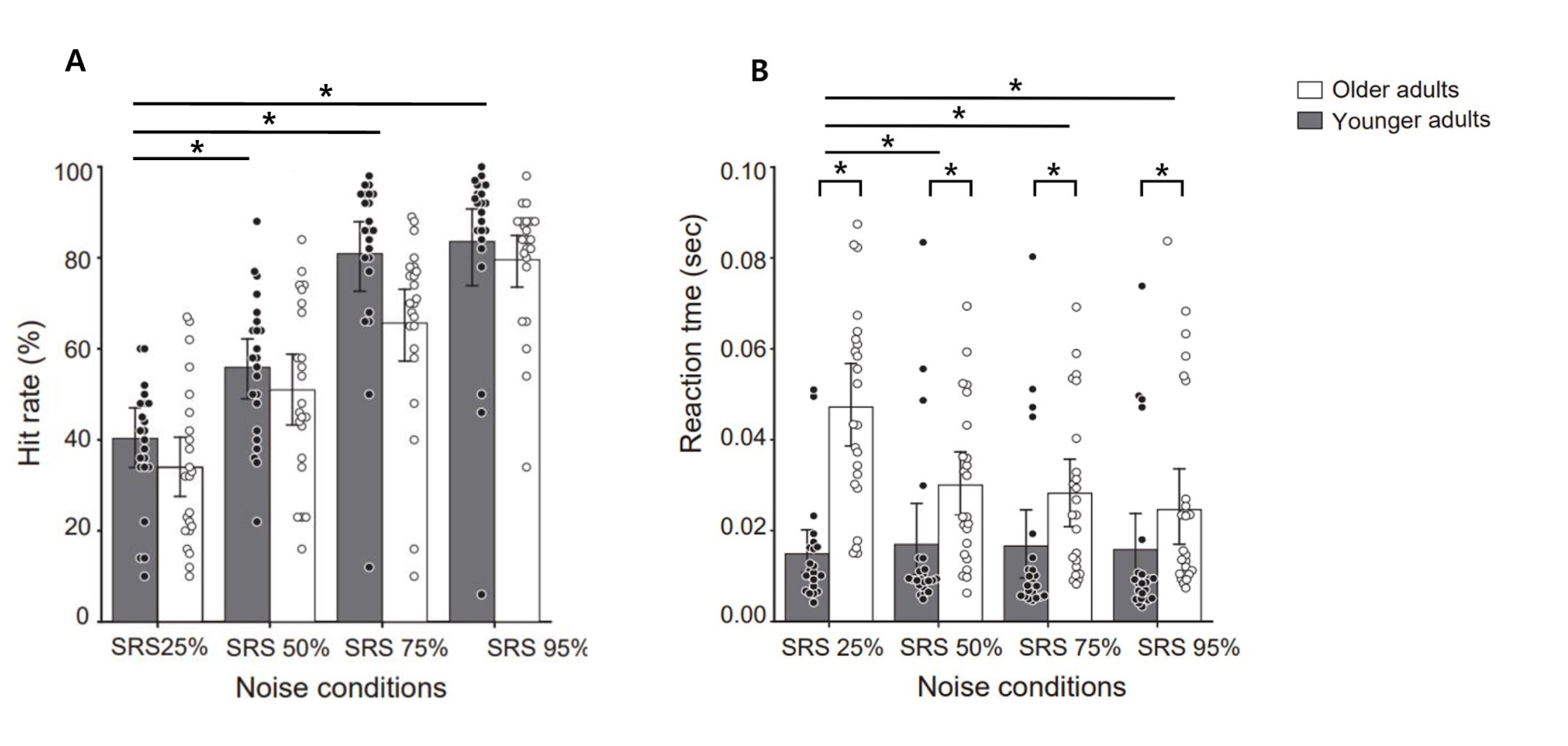
Behavioral result (hit rate and reaction time) under each noise level (SRS 25%, 50%, 75% and 95%). (A) The hit rate across different SRS noise conditions. The younger group is represented by dark bars, while the older group is represented by white bars. Each bar shows the mean hit rate, with individual participant data overlaid as dots. (B) The reaction times (in seconds) for both groups under the noise conditions, with the younger group data in dark bars and the older group in white bars. The reaction times for individual participants are shown as dots. Error bars represent the mean ± SEM. Black bars indicate *p < 0.05.

Regarding reaction time (Figure 2B), differences were observed between the younger and older groups. Both the SRS conditions (F_(3, 132)_ = 3.989, *p* = 0.0093, η^2^ = 0.28) and groups (F_(1, 44)_ = 19.43, *p* = 0.0001, η^2^ = 0.41) showed significant main effects, suggesting varied responses based on noise conditions and age group. A significant age group-by-SRS condition interaction (F_(3, 132)_ = 5.164, *p* = 0.00209, η^2^ = 0.32) indicated a differential performance decline in reaction time for both age groups. Post-hoc comparisons revealed a significant performance difference between the SRS 25% condition and the residual conditions (SRS 50%, 75%, and 95%) for the older group (*p* < 0.05). However, for the younger group, there was no significance across all SRS conditions. These results suggest that the older participants exhibited increased listening effort in severe noise level conditions.

### 3.2 Age-related differences in neural tracking of speech in noise

To investigate the differences in neural tracking of different frequency bands when perceiving speech in noise across age groups, we computed the speech envelope in the delta and theta bands based on the decoders trained while listening to clean story. The neural tracking data were subjected to a 2 (age group) × 4 (SRS conditions: SRS 25%, 50%, 75%, and 95%) mixed ANOVA, with the SRS condition as the within-subjects factor and age group as the between-subjects factor (Figure 3).

**Figure 3.**
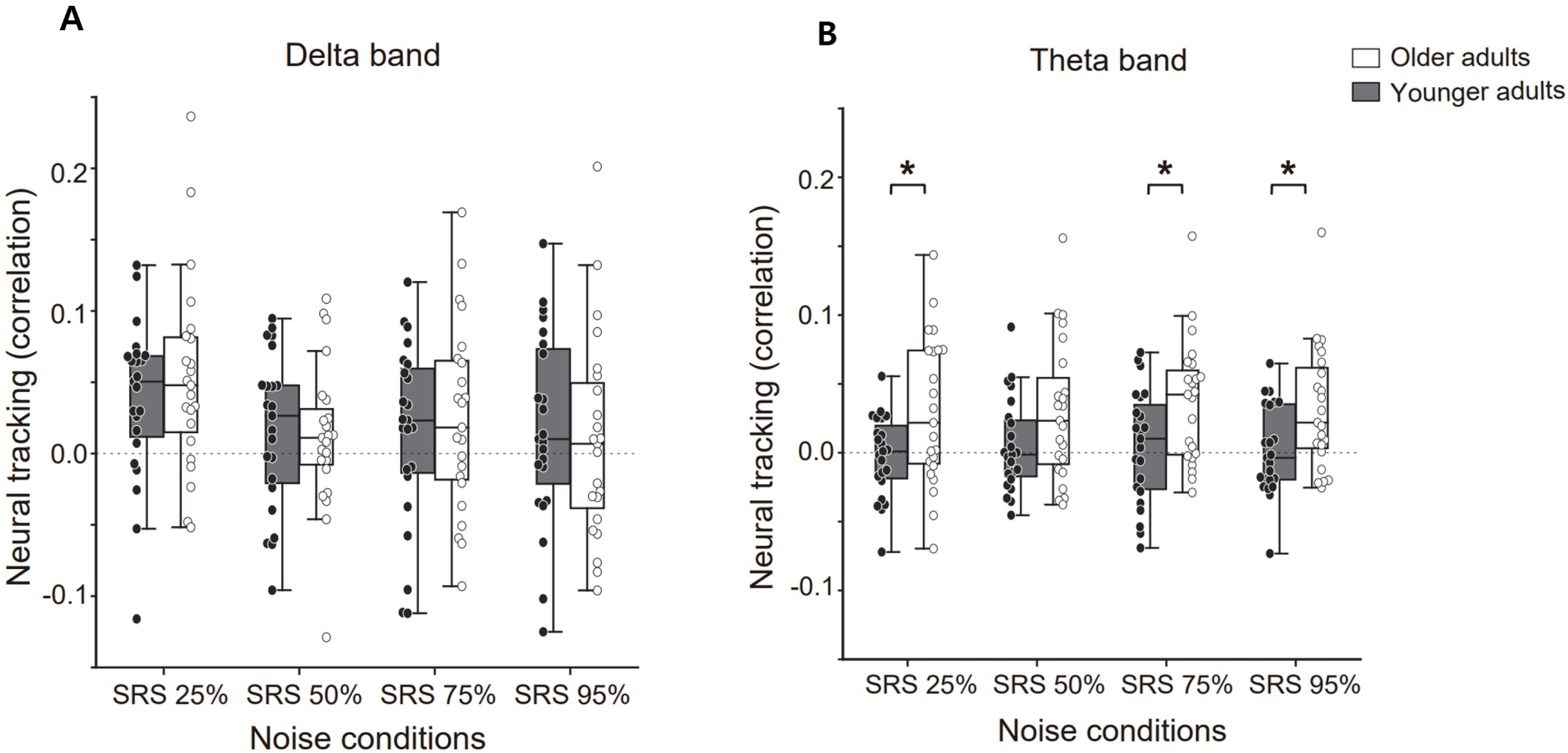
Neural tracking of speech in delta and theta bands across different noise conditions for older and younger groups. (A) The neural tracking within the delta frequency band across the SRS noise conditions (SRS 25%, 50%, 75%, and 95%). (B) For the theta frequency band. In both figures, the younger group is denoted by dark bars and the older group by white bars. Each dot represents an individual participant, and the central line on each box indicates the median value. Error bars represent the mean ± SEM. Black bar indicate *p < 0.05.

In the delta band, there was no significant main effect and interaction (Figure 3A). On the other hand, in the theta band, there was a main effect for the age group factor (F_(1, 44)_ = 11.06, *p* = 0.0017, η^2^ = 0.21), but no significance for the SRS conditions factor and interaction. Post-hoc comparisons demonstrated a significant difference in age group between all SRS conditions (*p* < 0.05) except the SRS 50%, but it also showed close statistical significance (*p* = 0.063) (Figure 3B). These findings suggest that the theta band might be more involved in speech recognition among older groups in noise conditions.

### 3.3 Correlations between cognitive function, age, and hearing ability in older adults (PTA and SRT)

To evaluate the interplay among cognition, age, and hearing ability, Pearson’s correlations were employed to determine the relationship between participants’ MMSE score and age, education level, as well as hearing ability. Figure 4 revealed that the MMSE score was significantly negatively correlated with age (*r* = −0.42, *p* = 0.046, Cohen’s d = 0.58) and significantly negatively correlated with the SRT (*r* = −0.46, *p* = 0.027, Cohen’s d = 0.54). However, there was no relationship with other factors (PTA, education level). This suggests a correlation between age-related cognitive decline and the deterioration of hearing sensitivity. The results showing the correlations between working memory task outcomes, age, and hearing ability (SRT and PTA) are presented in Supplementary Table 1. Please note that the significance levels displayed here have been adjusted for multiple comparisons.

**Figure 4.**
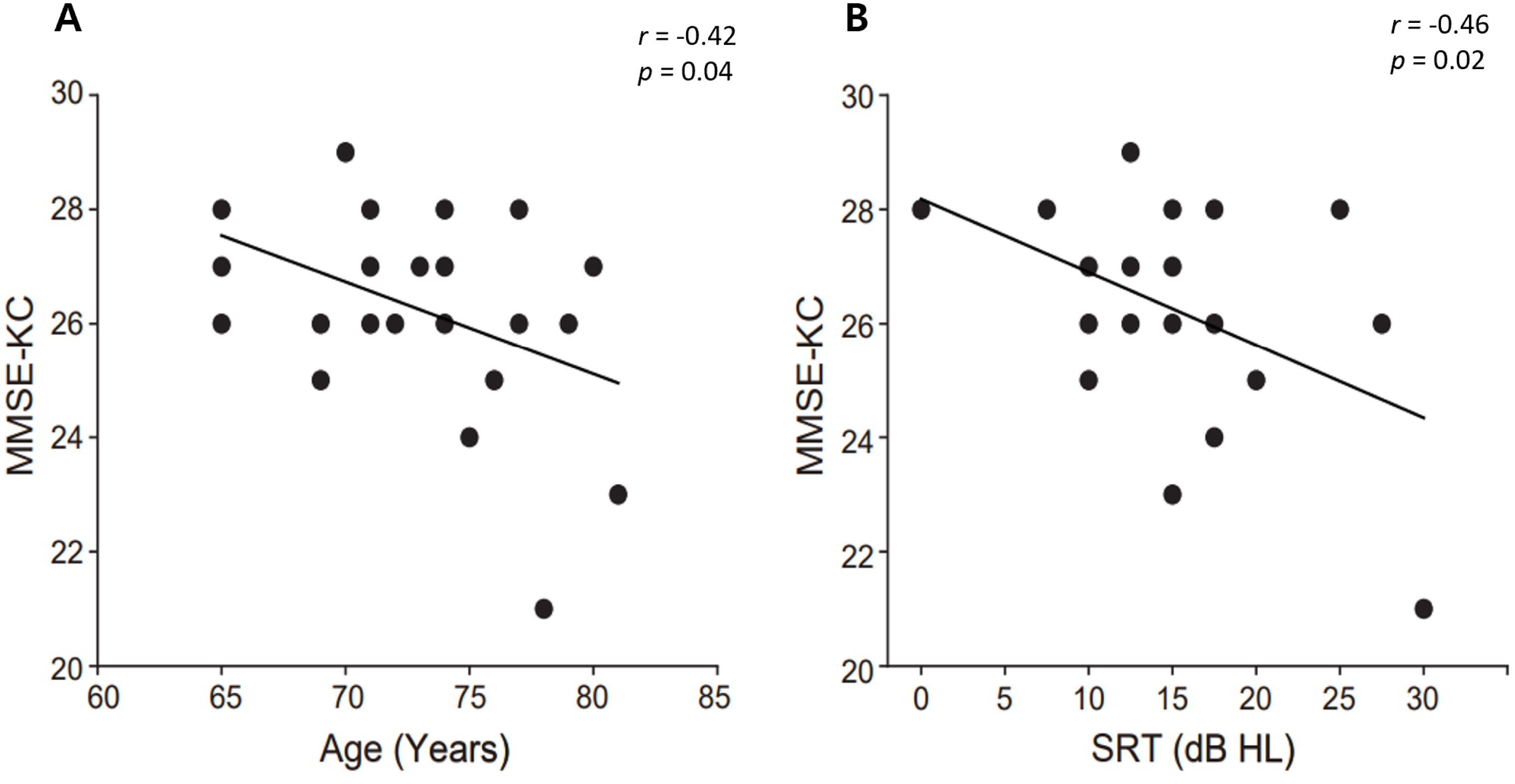
The correlation of MMSE-KC scores with age and the SRT. (A) The relationship between the MMSE-KC scores and age. Each dot represents an older adults score, with the trend line indicating the direction of the correlation. (B) The correlation between the MMSE-KC scores and the SRT. The trend line also indicates the correlation direction between the cognitive scores and hearing ability.

To identify the relationship between cognitive function and theta-band neural tracking at each noise level, a multiple linear regression analysis was conducted. The dependent variables were cognitive function measures (MMSE and working memory tests: word list memory, word list recall, word list recognition, and Constructional recall). Table 2 shows the results of the multiple regression analysis for each cognitive function test. The regression equation for word list recognition was significant, F(4, 18) = 3.427, *p* < .05, accounting for 43.2% of the variance (R^2^ = .432). Neural tracking at SRS 25% and SRS 75% significantly contributed to the predictions (*p* < .05). Except for word list recognition, other working memory test did not show significant contributions.

**Table 2.** Summary of the results of multiple regression analyses for each computerized cognitive function tests with predictors (n=23)

| Subtest Score | Predictor | Standardized beta coefficient | R square |
| --- | --- | --- | --- |
| MMSE | Neural tracking in SRS25 | -0.110 | 0.118 |
|  | Neural tracking in SRS50 | 0.282 |  |
|  | Neural tracking in SRS75 | 0.027 |  |
|  | Neural tracking in SRS95 | -0.432 |  |
| Word list memory | Neural tracking in SRS25 | 0.238 | 0.024 |
|  | Neural tracking in SRS50 | 0.089 |  |
|  | Neural tracking in SRS75 | -0.049 |  |
|  | Neural tracking in SRS95 | -0.056 |  |
| Word list recall | Neural tracking in SRS25 | -0.286 | 0.317 |
|  | Neural tracking in SRS50 | 0.418 |  |
|  | Neural tracking in SRS75 | -0.465 |  |
|  | Neural tracking in SRS95 | 0.107 |  |
| Word list recognition | Neural tracking in SRS25 | 0.683* | 0.432* |
|  | Neural tracking in SRS50 | 0.163 |  |
|  | Neural tracking in SRS75 | -0.997* |  |
|  | Neural tracking in SRS95 | 0.056 |  |
| Constructional recall | Neural tracking in SRS25 | 0.4470 | 0.086 |
|  | Neural tracking in SRS50 | 0.191 |  |
|  | Neural tracking in SRS75 | -0.281 |  |
|  | Neural tracking in SRS95 | -0.262 |  |

\* $p < 0.05$ , \*\* $p < 0.01$ SRS: Speech Recognition Scorer MMSE: Mini-Mental Status Examination-Korea

### 3.4 Correlation with working memory of older adults with different hearing abilities

Given the observed relationship between word list recognition and theta-band neural tracking in noise, we conducted an additional multiple regression analysis to test whether theta-band neural tracking was correlated with cognitive functions, particularly working memory based on hearing ability. Specifically, the two groups were divided based on the median value of their SRT levels, which was 15 dB HL. The group with thresholds higher than 15 dB HL was classified as the higher SRT group, while the group with thresholds lower than the median value was classified as the lower SRT group (Table 3). For the lower SRT group, a significant regression solution was obtained for constructional recall, F(4, 8) = 8.064, *p* < .05, accounting for 80.1% of the variance (R^2^ = .801). This is also indicated by a positive correlation for neural tracking at SRS 25% and SRS 50%, and a negative correlation for neural tracking at SRS 95% as significant predictors (*p* < .05). On the other hand, for the higher SRT group, the regression equation for word list recognition approached significance, F(4, 5) = 4.632, *p* = .052, accounting for 78.7% of the variance (R^2^ = .787), with a positive correlation for neural tracking at SRS 25% and a negative correlation for neural tracking at SRS 75% as significant predictors (*p* < .05).

**Table 3.** Summary of the results of multiple regression analyses for working memory tests with predictors (lower SRT group=13, higher SRT group=10)

| Group | Subtest Score | Predictor | Standardized beta coefficient | R square |
| --- | --- | --- | --- | --- |
| lower SRT group | Word list memory | Neural tracking in SRS25 | 0.302 | 0.419 |
|  |  | Neural tracking in SRS50 | 0.384 |  |
|  |  | Neural tracking in SRS75 | 0.667 |  |
|  |  | Neural tracking in SRS95 | -0.801 |  |
|  | Word list recall | Neural tracking in SRS25 | -1.556 | 0.507 |
|  |  | Neural tracking in SRS50 | -0.740 |  |
|  |  | Neural tracking in SRS75 | -0.937 |  |
|  |  | Neural tracking in SRS95 | 1.645 |  |
|  | Word list recognition | Neural tracking in SRS25 | 0.329 | 0.270 |
|  |  | Neural tracking in SRS50 | -0.070 |  |
|  |  | Neural tracking in SRS75 | -0.132 |  |
|  |  | Neural tracking in SRS95 | -0.446 |  |
|  | Constructional recall | Neural tracking in SRS25 | 0.958** | 0.801* |
|  |  | Neural tracking in SRS50 | 0.691** |  |
|  |  | Neural tracking in SRS75 | -0.077 |  |
|  |  | Neural tracking in SRS95 | -0.713* |  |
| higher SRT group | Word list memory | Neural tracking in SRS25 | 0.032 | 0.114 |
|  |  | Neural tracking in SRS50 | 0.169 |  |
|  |  | Neural tracking in SRS75 | -0.558 |  |
|  |  | Neural tracking in SRS95 | 0.236 |  |
|  | Word list recall | Neural tracking in SRS25 | 0.103 | 0.540 |
|  |  | Neural tracking in SRS50 | 1.426 |  |
|  |  | Neural tracking in SRS75 | -1.015 |  |
|  |  | Neural tracking in SRS95 | -0.055 |  |
|  | Word list recognition | Neural tracking in SRS25 | 0.888* | 0.787* |
|  |  | Neural tracking in SRS50 | 0.715 |  |
|  |  | Neural tracking in SRS75 | -1.536* |  |
|  |  | Neural tracking in SRS95 | -0.087 |  |
|  | Constructional recall | Neural tracking in SRS25 | -0.203 | 0.453 |
|  |  | Neural tracking in SRS50 | -0.638 |  |
|  |  | Neural tracking in SRS75 | 0.074 |  |
|  |  | Neural tracking in SRS95 | 0.004 |  |

\*<0.05, \*\*p<0.01 SRT: Speech Recognition Threshold

## 4. Discussion

### 4.1 Age-related changes in speech perception in noise

In the current study, we investigated the effect of age on speech recognition performance among individuals with normal hearing. It was observed that as the SRS decreased, the hit rate for the older group decreased, while the reaction time increased (Figure 2). These findings are consistent with previous studies, which show that older listeners are more adversely influenced by background noise compared to younger listeners (Wingfield et al., 1985; Yanz and Anderson, 1984). Yanz and Anderson (1984) reported a notable increase in the precision of word recognition among the younger participants when the SNR was adjusted from 0 to 5 dB. However, older listeners showed no significant change under all SNR conditions. Furthermore, Wingfield et al. (1985) suggested that the diminished ability of older adults to successfully perform speech recognition tasks might be linked to their less effective use of contextual information. Also, when younger and older adults listen to speech at the SNR 0 dB, older people often find it harder to understand, need more effort (Houben et al., 2013), and take longer to respond (Hultsch et al., 2002; Salthouse, 2010). As people age, the difficulty in recognizing speech in noise is caused not only by a decline in the peripheral auditory system but also by a decline in central auditory function (Dobreva et al., 2011; Helfer and Wilber, 1990). Several studies show that deficits in central auditory function could make it more difficult to distinguish target sounds from background noise as the noise level increases (Ben-David et al., 2012; Snyder and Alain, 2005). Previous studies have also reported that cognitive overload caused by external variables such as background noise during listening tasks can lead to increased response time on the task or abandonment of the task (Brungart et al., 2020; Wu et al., 2016). These findings support our results, which show that older adults, compared to younger adults, have slower response times as noise levels increase. This suggests that older adults experienced greater listening effort under severe noise conditions.

### 4.2 Neural tracking changes across aging

We conducted further investigations to understand how neural tracking during speech recognition in noise differs according to age. Our results showed that delta-band neural tracking was not differ between younger and older listeners in any of the SRS conditions. However, theta-band neural tracking was greater for older listeners than younger listeners. Our findings, consistent with Kurthen et al. (2022) highlight age-related adaptations in neural processing of speech in challenging auditory conditions. While delta-band neural tracking did not differ significantly between younger and older adults in our study, older listeners showed greater theta-band neural tracking compared to younger listeners. They demonstrated that this enhanced theta-band tracking in older adults was positively associated with understanding interrupted speech, suggesting a compensatory mechanism for processing degraded auditory input. Similarly, our findings indicate that older adults might use increased theta-band tracking to support speech recognition in noise, potentially compensating for age-related declines in auditory processing.

Previous studies have shown that theta oscillations reflect neural sensitivity to the slow-rate envelope in the auditory cortex (Howard and Poeppel, 2010; Luo and Poeppel, 2007; Rosen, 1992), which plays a critical role in speech perception in noise (Arai et al., 1999; Drullman et al., 1994; Shannon et al., 1995). Specifically, research on neural tracking in the theta frequency band has demonstrated that it is a significant predictor of perceived speech clarity, primarily reflecting the acoustics of the stimulus. In contrast, neural tracking in the delta band has been found to predict speech comprehension, suggesting an association with higher-level linguistic aspects of speech, such as syntax and semantics. This indicates that while theta band tracking is linked to the physical properties of sound, delta band tracking relates to the understanding of speech content (Di Liberto et al., 2015; Ding et al., 2014; Donhauser and Baillet, 2020; Etard and Reichenbach, 2019). Furthermore, the enhanced neural tracking observed in older adults and those with hearing impairment is thought to result from central compensatory mechanisms that increase cortical activity to compensate for the degraded peripheral input (Chambers et al., 2016; Decruy et al., 2019; Schmitt et al., 2022). In this context, the observed increase in theta band neural tracking in our study may reflect neural compensatory mechanisms in response to the diminished clarity of phoneme and syllable boundaries due to the presence of background noise.

### 4.3 Relation between neural tracking and working memory in older adults

The correlation analysis on MMSE and hearing ability revealed that general cognitive function (MMSE) showed a negative correlation with SRT and age. This finding is consistent with previous studies that have demonstrated a significant relationship between MMSE scores and demographic variables such as age and education in the general population (Brayne and Calloway, 1990; Pliatsikas et al., 2019; Weiss et al., 1995; Ylikoski et al., 1992). To extend the correlation analysis results, we also performed a linear regression analysis between the scores of working memory test in CERAD-K (word list recall, word list memory, word list recognition, and constructional recall) and theta-band neural tracking of the speech envelope across different noise levels. With regard to working memory, we found that theta-band neural tracking could significantly predict the performance of word list recognition (Table 2).

Furthermore, we explored the relation between cognitive function and speech perception by categorizing the participants into two groups according to their ability to recognize speech in noise. Specifically, the two groups were divided based on the median values of their SRT levels. Our study observed that older adults with lower SRT and higher SRT group showed different correlation patterns in theta-band neural tracking across different working memory tasks (word list recognition and constructional recall).

In the lower SRT group, the regression analysis results indicate that SRS 25% and SRS 50% have a significant positive effect on constructional recall. In particular, SRS 25% has the highest standardized coefficient (β = .958) and t-value (4.480), making it the strongest predictor of visual working memory score. Conversely, in the higher SRT group, the regression analysis results showed that the SRS 25% noise condition has a significant positive effect on word recognition ability (Table 3).

In higher SRT group, the positive relationship with word list recognitions, which related the verbal working memory, can potentially be explained by the information degradation hypothesis. According to this hypothesis (Wayne and Johnsrude, 2015), older adults may increasingly rely on cognitive resources to compensate for sensory deficits. This reliance, in turn, can deplete the resources available for other cognitive tasks, resulting in a comparative decline in cognitive performance relative to younger adults. In this regard, the information degradation hypothesis may explain why the higher SRT group exhibited an increased envelope correlation at higher word list recognition specially in SRS 25% which is severe noise conditions. Individuals in the higher SRT group, indicating hearing deficits, may have exerted more effort in listening tasks with a greater reliance on verbal working memory. Indeed, previous studies have also confirmed that sensory deficits are associated with increased listening effort and greater demands on working memory resources (Amichetti et al., 2013; McCoy et al., 2005; Wingfield and Tun, 2007). Similarly, although age groups differ, several studies have found that children with cochlear implants (CIs) perform worse than normal hearing peers in complex verbal working memory (Pisoni and Cleary, 2003; Kronenberger et al., 2013; Watson et al., 2007).

In contrast, among those with a lower SRT, a positive correlation was observed between neural tracking and the constructional recall, which reflects the visual working memory. This finding could align with prior research indicating increased visual cortex activity in the elderly when trying to perceive speech in noise (Kuchinsky et al., 2012; Peelle et al., 2010) but further study is necessary to understand the relationship between visual-working memory and neural tracking of speech in noise for older adults.

Additionally, both SRT groups showed a negative correlation in the SRS 75% and SRS 95% conditions for word list recognition and constructional recall. This observation could be explained by the Ease of Language Understanding (ELU) model. According to the ELU model, when listening conditions are optimal, speech input is quickly and automatically matched with phonological representations stored in semantic long-term memory, allowing access to the lexicon. This process does not place a burden on working memory, which involves the ability to simultaneously process and store phonological and semantic information and to draw inferences for both predictive and reflective purposes (Baddeley, 2012; Ronnberg et al., 2013; Ronnberg et al., 2019). Therefore, the SRS 75% and 95% conditions, which have less noise compared to the SRS 25% and 50% conditions, exert less demand on working memory processing.

### 4.4 Limitations and future work

In this study, we investigated the relationship between cognitive function and neural envelope tracking of speech in noise. The regression analysis revealed that the SRT groups exhibit different patterns of correlation with the working memory task. However, the regression results alone are not sufficient to fully explain how working memory influences neural envelope tracking of speech in noise. One of the main limitations of our study is the difficulty in directly explaining the relationship between auditory cognitive ability in processing speech in noise, cognitive function, and language skills. Although our study provides valuable insights into the role of working memory in speech understanding, we did not simultaneously assess other cognitive factors, such as attention, linguistic skills, which have been reported to impact speech perception in noise (Kilman et al., 2014; Magimairaj et al., 2021). As a result, our findings, while significant, may not fully capture the complex interplay between cognitive and linguistic skills and neural envelope tracking of speech in noise. To build upon our current findings and gain a more comprehensive understanding of these relationships, future research should aim to incorporate a more comprehensive battery of tests, assessing a broader range of cognitive functions and language ability alongside EEG data collection. This approach would enable a more in-depth understanding of how these factors contribute to speech perception in challenging listening environments and their direct influence on neural envelope tracking. Also, hearing loss or lack in temporal spectral encoding and other sensory inputs can affect cognitive functions (Wong et al., 2009). Previous studies have shown that patients with deficits in auditory attention and memory have difficulty in understanding speech in noisy environments (Frisina and Frisina, 1997; Tun et al., 2002). Since our study was conducted on older adults with normal hearing, it is also difficult to completely separate and interpret pure auditory ability from cognitive function. Additionally, mild cognitive impairment (MCI) can affect the neural processing of speech in noise, but our study did not adequately capture the full spectrum of cognitive impairments. These limitations mean that we could not fully address the hypothesis concerning the interplay between cognitive functions and neural processing of speech in noise. Future research should include a broader cohort of older adults with varying levels of cognitive impairments and hearing abilities, encompassing those with normal hearing and those with hearing deficits. This would allow for a more thorough investigation into how different cognitive abilities interact with auditory processing, potentially providing a clearer understanding of the underlying mechanisms.

Another limitation of our study is that it is difficult to clearly distinguish and explain the processing of speech in noise as either pure sensory processing or cognitive processing. Perceiving speech in noise involves a complex interplay of auditory and cognitive processes. Both bottom-up processing, which involves the direct analysis of acoustic signals, and top-down processing, which uses context and prior knowledge, are crucial for speech in noise processing (Zekveld et al., 2006). In this study, we measured sentence recall under noise conditions by having older adults with normal hearing listen to sentences in noisy environments. Additionally, neural envelope tracking was measured by correlating the original envelope of the matrix sentences with the envelope reconstructed from the EEG responses. In this regard, this study liikely involved the top-down processing, as the task required focusing on target sentences in noise and recalling the perceived sentence. In contrast, a recent study by Mai and Howell (2023) investigated how early-stage phase-locked neural activities, specifically the frequency-following response (FFR) and theta-band phase-locking values (θ-PLV), contribute to speech-in-noise (SiN) perception across different ages and levels of hearing loss. Their study provided evidence of distinct early-stage neural mechanisms through which aging and hearing loss affect speech-in-noise perception. Based on prior research designs, future studies should design additional experiments that use vowel acoustic stimuli to investigate early-stage neural activity, considering both pure sensory processing and processes involving cognitive functions.

## 5. Conclusion

In conclusion, we aimed to investigate the relationship between hearing ability, cognitive function, and age. We explored the effect of age and hearing ability on neural and cognitive factors in speech recognition in noisy environments through neural tracking analysis, with a particular focus on the roles of delta and theta cortical oscillations. Our hypothesis was that age-related differences in neural tracking in noise would exist due to differing mechanisms for speech recognition in noise, and that these differences would be related to working memory. Our results showed behavioral differences in speech recognition based on age and revealed significant differences in theta-band neural tracking between age groups. We also found that theta-band neural tracking is particularly associated with working memory, specifically with word list recognition. Lastly, our study found that older adults with lower and higher SRT levels exhibited different correlation patterns in theta-band neural tracking across various working memory tasks. Taken together, these findings suggest that mechanisms of speech perception in noise could differ according to age, and we indirectly identified a connection between speech perception in noise and working memory. In future studies, we aim to clarify the causal relationship between cognitive function and hearing by addressing the limitations of this research.

